# Efficacy of anti-microbial gel vapours against aerosolised coronavirus, bacteria, and fungi

**DOI:** 10.1101/2021.10.27.466182

**Authors:** Parthasarathi Kalaiselvan, Muhammad Yasir, Mark Willcox, Ajay Kumar Vijay

## Abstract

**Background:** The urban population spends up to 90% of their time indoors. The indoor environment harbours a diverse microbial population including viruses, bacteria, and fungi. Pathogens present in the indoor environment can be transmitted to humans through aerosols.

**Aim:** This study evaluated the efficacy of an antimicrobial gel containing a mix of essential oils against aerosols of bacteria, fungi, and coronavirus.

**Methods:** The antimicrobial gel was allowed to vapourize inside a glass chamber for 10 or 20 minutes. Microbial aerosols of *Escerichia coli, Aspergillus flavus* spores or murine hepatitis virus MHV 1, a surrogate of SARS CoV-2 was passed through the gel vapours and then collected on a 6-stage Andersen sampler. The number of viable microbes present in the aerosols collected in the different stages were enumerated and compared to number of viable microbes in control microbial aerosols that were not exposed to the gel vapours.

**Results:** Vaporizing the antimicrobial gel for 10 and 20 minutes resulted in a 48% (p = 0.002 Vs. control) and 53% (p = 0.001 Vs. control) reduction in the number of MHV-1 in the aerosols, respectively. The antimicrobial gel vaporised for 10 minutes, reduced the number of viable *E. coli* by 51% (p = 0.032 Vs. control) and *Aspergillus flavus* spores by 72% (p=0.008 Vs. control) in the aerosols.

**Conclusions:** The antimicrobial gel may be able to reduce aerosol transmission of microbes.

## INTRODUCTION

The majority of the urban population spend up to 90% of their time indoors [1, 2]. The indoor environment harbours a diverse microbial population including viruses, bacteria, fungi and protozoa [3-6] that is referred to as the indoor microbiome. A major component of indoor microbiome are endogenous microbes shed by human and animal occupants with a minor constituent being the transient microbiota of external environment transported inside [7]. Additional sources that can contribute to the indoor microbiome include water from indoor plumbing such as toilets and showers, soil, heating, and ventilation systems such as air-conditioning systems [4, 8-12]

Human exposure to the indoor microbiome has been recognised as a factor for the development of respiratory diseases and allergies. Pathogens present in the indoor microbiome can be transmitted to humans either through aerosols or from contaminated surfaces. Key pathogens that are transmitted through aerosols include the bacteria *Staphylococcus aureus* [13], *Mycobacterium tuberculosis* [14], the fungus *Aspergillus fumigatus* [15], and the viruses Influenza virus, Ebola and SARS-CoV [16]. While there was considerable speculation regarding the aerosol transmission of the SARS-CoV-2 virus [17, 18], current data confirms aerosol transmission of this virus [19, 20].

Hospitals and food industries use UV-C irradiation, plasma air ionization and fumigation with disinfectants to reduce air borne pathogens in the indoor air [21, 22]. However, these strategies are expensive and may not be suitable in domestic settings. Portable indoor air cleaners/purifiers with HEPA filters are effective in reducing the microbial concentration in aerosols including SARS-CoV-2 in classrooms, offices, and hospitals [23-25]. Vapours of essential oils have good antimicrobial activity against respiratory pathogens [26, 27] and offer an alternative strategy for disinfecting the indoor air [28-30]. The vapours when dispersed in the air can significantly reduce the microbial levels indoors [31-33]. In this study we evaluated the antimicrobial efficacy of an antimicrobial gel containing a proprietary mix of essential oils for its activity against pathogenic bacteria, fungi, and a coronavirus surrogate of SARS-CoV-2.

## MATERIALS AND METHODS

### Microorganisms and their preparation

The mouse hepatitis virus (MHV-1) ATCC/VR261 is an enveloped single-strand RNA virus and an accepted surrogate of the SAR-CoV-2 virus. Viral stock was prepared by growing in A9 mouse fibroblast cells (ATCC/CCL 1.4) in Dulbecco’s minimum essential medium (DMEM, Thermofisher, Macquarie Park, NSW, Australia) containing 10% foetal bovine serum (FBS; Thermofisher), 100 μg/ml streptomycin sulphate and 100 I.U. penicillin G, (Thermofisher). Viral titres (1.0 × 10^5^ to 1.0 × 10^6^ plaque forming units (PFU)/mL) were determined by plaque assay as described below.

*Escherichia coli* K12 (ATCC 10798) was grown overnight in tryptic soy broth (TSB; BD, Sydney, NSW, Australia) to mid-log phase. Following incubation, bacterial cells were collected by centrifuging and were washed once with phosphate buffer saline (PBS; NaCl 8 g/L, KCl 0.2g/L, Na_2_HPO_4_ 1.15 g/L, KH_2_PO_4_ 0.2 g/L, pH 7.4). Following washing, cells were re-suspended in PBS and the concentration adjusted spectrophotometrically to an optical density of 0.1 at 660 nm which yielded 1.0 × 10^8^ colony forming units (CFU/mL) upon retrospective agar plate counts, then further serially diluted to a final concentration of 1.0 × 10^4^ CFU/mL.

The spores of *Aspergillus flavus* ATCC 9643 were produced by growth on Sabouraud dextrose agar (SDA; Thermofisher) for 10 days at 25°C. The fungal growth was suspended in sterile deionized water and filtered through sterile 70 μm filters to remove hyphal fragments. Spores were resuspended in sterile deionized water and their concentration adjusted spectrophotometrically to an optical density of 0.2 at 660 nm which yielded 1.0 × 10^6^ CFU/mL, which were then serially diluted to a final concentration of 1.0 × 10^4^ CFU/mL.

### Antimicrobial Gel

The antimicrobial gel (Mould Gone, SAN-AIR, West Gosford, NSW, Australia) was supplied in sealed containers. The gel formulation is proprietary but includes monoterpenoids, diterpenoids and sesquiterpenoids found in essential oils of plants from the *Melaleuca* genus (e.g., 1,8 cineole, thymol, alpha-pinene, caryophyllene) with an average dose of 0.0005% (v/v) when vapourised according to the manufacturer.

### Activity of the antimicrobial gel against coronavirus in solution

In order to demonstrate that the gel had antiviral activity, the first experiments incubated aliquots of the gel directly with viral particles in suspension. Cells of MHV-1 (1.0 × 10^5^ (PFU)/mL) were incubated with 25 mg or 50 mg of the antimicrobial gel in DMEM at ambient temperature for 0.5 or 2 hours. Following incubation, the DMEM was removed, diluted in 20% (w/v) bovine serum albumin (BSA; Sigma-Aldrich, Castle Hill, NSW, Australia) prepared in PBS and incubated for 10-15 minutes to neutralize the antimicrobial agents released from the gel. Thereafter, 100 μL aliquots were diluted ten-fold (in 20% BSA) and inoculated into the wells of 12 well plates containing A9 cells and incubated for 1 hour at 37 °C in the presence of 5% (v/v) CO_2_. The plates were gently rocked once every 15 minutes to prevent the cells from drying out. After incubation, an overlay media containing a 50:50 mix of 2% (w/v) agar (Sigma-Aldrich) and DMEM was added to each well and further incubated for 72 hours. Following incubation, the cells were fixed with 4% (v/v) formaldehyde (Sigma-Aldrich) for 2 – 3 hours, the agar overlay removed, and the number of viral particles (PFUs) visualized after staining with 1% (w/v) crystal violet (Sigma-Aldrich). Controls were the viral inoculum incubated in DMEM or PBS without the antimicrobial gel. The percentage reduction in PFU for each quantity of the gel compared to the negative control (PBS) was calculated.

### Activity of the gel as vapours against viral aerosols

As there are no standard assays for examining the effects of vapourised or aerosolised disinfectants on aerosols of microbial cells, a new assay was devised. A bacterial filtration efficiency (BFE) test rig (CH Technologies, Westwood, NJ, USA) was used to produce viral aerosols (Figure 1). The antimicrobial gel (10 g) was removed from its container and allowed to vaporise into the glass aerosol chamber for 10 minutes or 20 minutes prior to the introduction of the virus. The viral inoculum (50 μL; 1.0 × 10^6^ PFU/mL) was aerosolized using a continuous drive syringe pump through a nebulizer with an airflow of 28.3 L/min for one minute and allowed to interact with vapours of the antimicrobial gel as they passed through the glass tube. The size of the aerosols produced was approximately 3.0 ± 0.3 μm and these travelled through the glass aerosol chamber into an Anderson sieve sampler and were collected by flowing past 2% (w/v) agar plates. The largest (7 μm) sized aerosols were captured on the agar plate at the top of the Anderson sieve and the smallest (0.65 μm) on the agar plate at the bottom of the device. After one minute, the airflow was stopped to cease aerosol generation, and the vacuum pump was run for further one minute to collect any residual aerosols from the glass chamber. Following this, agar plates were flooded with 1.5 ml of either 20% BSA in DMEM (neutralised samples) or DMEM alone (non-neutralised samples), and viruses were carefully removed using a sterile cell scrapper. Aliquots (100 μL) from each plate were placed in duplicate on A9 cells in 12-well cell culture plates to culture any infectious viruses. The culture conditions were as described above. Control runs were performed at the beginning of each experiment prior to the addition of the gel in the glass aerosol chamber to collect infectious viruses so that any reduction in the number of infectious viruses could be calculated as a percentage of this control. Test and control runs were conducted in duplicate and repeated twice.

**Figure 1:**
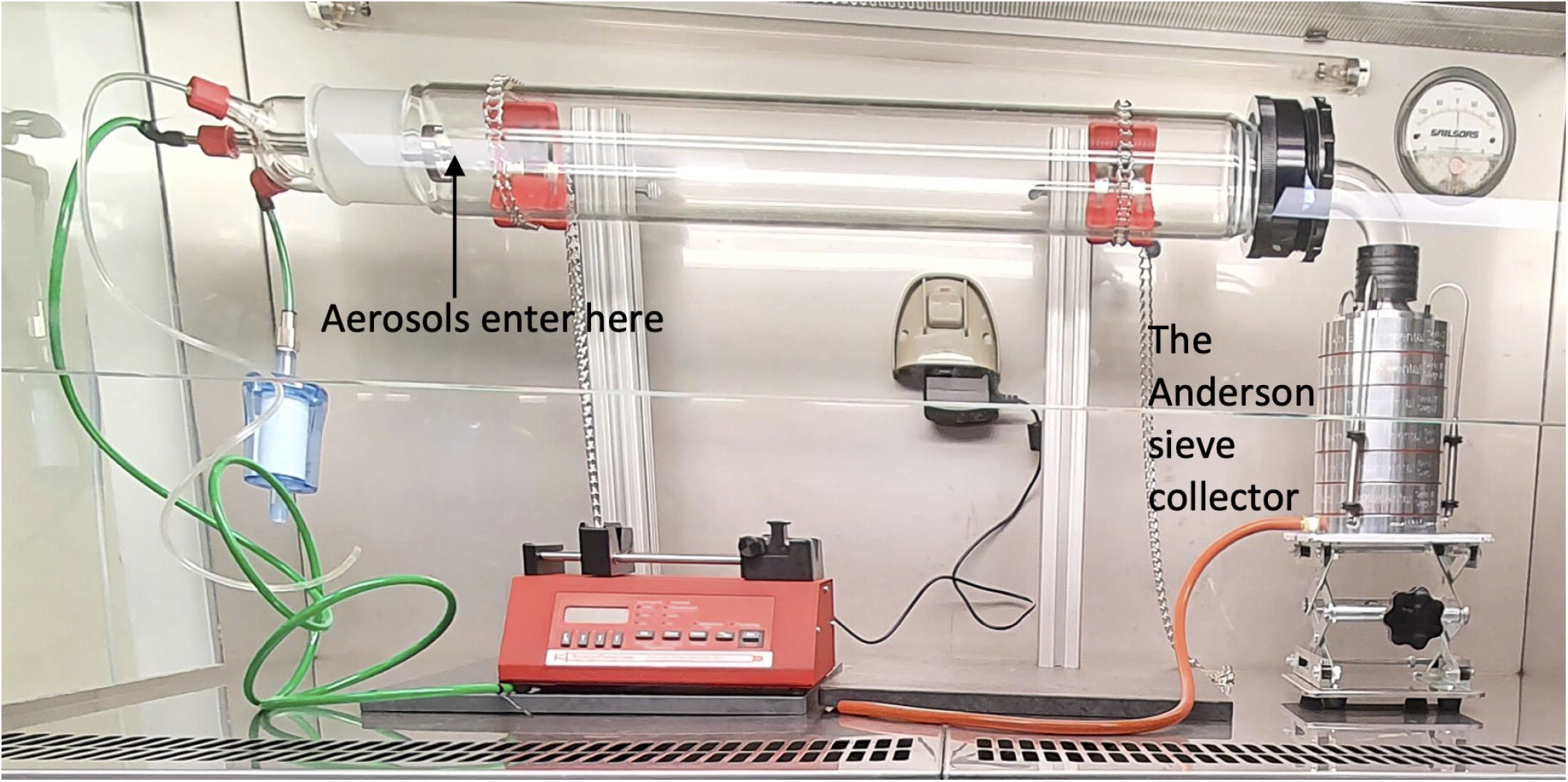
The Bacterial Filtration Efficiency Rig containing an Anderson sieve sampler. Aerosols of 3.0 ± 0.3 μm on average of viruses, bacteria or fungal spores were produced in the glass chamber.

### Activity gel as vapours against bacterial and fungal spore aerosols

The anti-bacterial activity of the gel vapourised for 10 minutes against *E. coli* and its sporicidal activity against *A. flavus* spores was determined using a similar method as described for MHV-1, except using 50 μL of *E. coli* or *A. flavus* spores (1 × 10^4^ CFU/mL). Bacteria were collected on agar plates composed of tryptic soy agar (TSA; BD, Macquarie Park, NSW, Australia) alone or containing TSA and the neutralizers Tween® 80 (5 g/L) and lecithin (7 g/L). Fungal spores were collected on SDA plates alone or containing the same neutralizers. The numbers of viable cells from each of the 6 plates in the Anderson sieve collector were enumerated following incubation at 37 ºC for 24 hours for bacteria and at 25 ºC for 72 hours for fungal spores. Control runs were conducted prior to the addition of the gel in the glass aerosol chamber to collect viable bacteria, and fungal spores. Test and control runs were performed in duplicate and repeated twice. The percentage of cells remaining viable after passage through the gel vapours was calculated by comparing numbers in the absence (control) and presence (test) of the gel vapours.

### Statistical analysis

Statistical analyses were performed using GraphPad Prism 7.04 software (GraphPad Software, La Jolla, CA, USA). The concentration and time dependent effect of the antimicrobial gel in solution was determined using two-way ANOVA. The effect of antimicrobial gel vapours at single time points on different aerosols sizes and overall percentage (%) reduction was assessed using Welch’s t-test and one-way ANOVA with Tukey’s test respectively. Statistical significance was set as P□<□0.05.

## RESULTS

### Activity of the gel in solution against coronavirus

The antimicrobial gel when incubated in DMEM with the coronavirus reduced the numbers of infectious MHV-1 in solution in a dose dependent manner. The greatest quantity (50 mg) of the gel reduced the infectivity of the coronavirus by >99.99% (no viral cells were cultured) within 30 minutes of incubation compared to control (p < 0.001; Table I). The smaller quantity (25 mg) of the gel reduced the numbers of coronavirus by 98.6% after 30 minutes of incubation compared to control (p < 0.001).

**Table I:** Effect of different concentrations of antimicrobial gel against coronavirus MHV-1 in solution at different time points.

| Sample | Amount of gel (mg) | Incubation time (minutes) | Number of plaques (PFU/mL) | Log<sub>10</sub> reduction | % Reduction | P-value |
| --- | --- | --- | --- | --- | --- | --- |
| Control | - | - | 26750 ± 1658 | - | - |  |
| Antimicrobial gel | 25 | 30 | 375 ± 35 | 1.9 | 98.60 | <0.001 |
|  | 25 | 120 | 225 ± 177 | 2.1 | 99.16 | < 0.001 |
|  | 50 | 30 | 0 ± 0 | 4.4 | 99.99 | < 0.001 |
|  | 50 | 120 | 0 ± 0 | 4.4 | 99.99 | <0.001 |

### Activity of the gel as vapours against viral aerosols

The antimicrobial gel vapours were active against MHV-1 aerosols. The majority of the viral particles travelled in aerosols of 3.30 to 0.65 μm in the absence of the gel (Figure 2A) with most viral particles travelling in the 2.10 and 1.10 μm aerosols (Figure 2A). After allowing the gel to vaporize in the chamber for 10 minutes, the numbers of viral particles that were able to infect the mouse cells were reduced for most aerosol sizes, with a significant reduction of 67% in the 1.10 μm aerosol (p = 0.011; Figure 2A). A slightly greater reduction of 78% was produced in the 2.10 μm aerosols compared to the controls when the gel was allowed to vaporize for 20 minutes (p = 0.011; Figure 2B). Overall, exposure of MHV-1 aerosols to the antimicrobial gel vapours (vaporized for 10 minutes) resulted a significant 48% reduction of all the aerosols sizes compared to untreated control (p = 0.002; Table II). Allowing the antimicrobial gel to vaporize for 20 minutes, resulted in a 53% reduction in the number of viable aerosolized viral particles of all sizes compared to control (p = 0.001; Table II). Following neutralization with 20% BSA, the activity of the antimicrobial gel was slightly but not significantly (p = 0.078; Table II) reduced, resulting in a 33% reduction in the viability of viral aerosols compared to control (p = 0.001; Table II).

**Figure 2:**
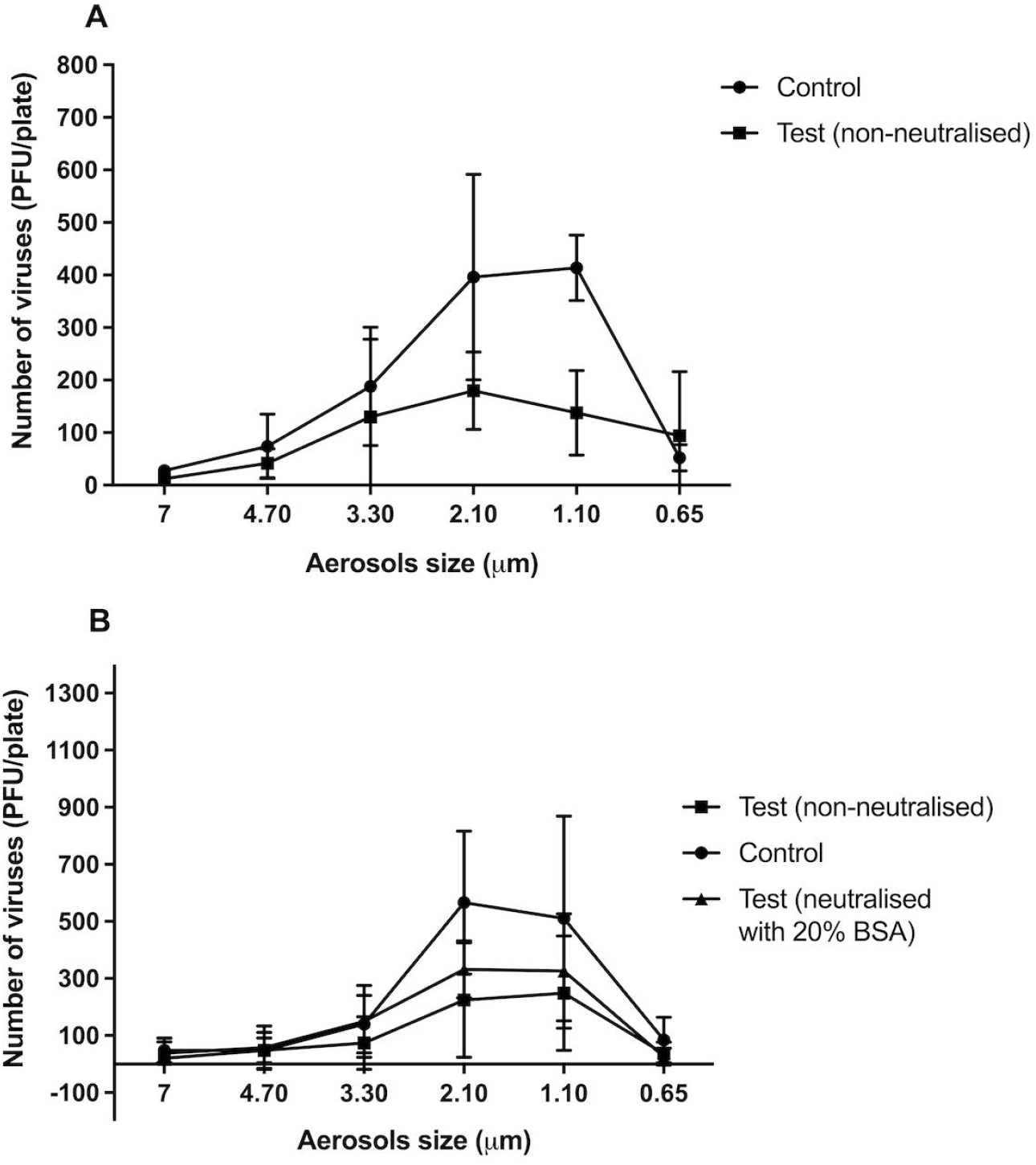
Number of murine hepatitis virus (MHV-1) recovered from different aerosol sizes with or without neutralization of the gel vaporized for 10 minutes (A) or 20 minutes (B). The antimicrobial gel significantly reduced the ability of viral aerosols to infected A9 cells in aerosols sizes of 2.10 and 1.10 μm compared to untreated control (p ≤ 0.001). Data points represent the mean (± 95% confidence interval) of three independent experiments.

**Table II:**
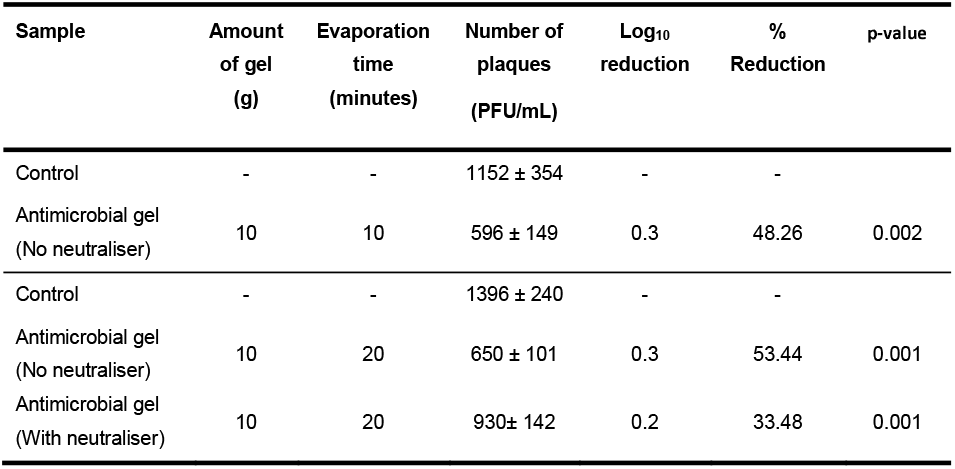
The ability of antimicrobial gel vapourised for 10 and 20 minutes to reduce the numbers of aerosolised coronavirus MHV-1.

### Activity against aerosols of bacteria or fungal spores

The antimicrobial gel in vaporized form was active against aerosols of *E. coli*. In the absence of antimicrobial gel, this bacterium mostly travelled in aerosol particle sizes between 3.30 to 1.10 μm (Figure 3). Overall, the antimicrobial gel produced a reduction in bacterial viability of 29% (p = 0.018) when neutralised during bacterial growth and 51% (p = 0.032) when not neutralised during bacterial growth (Table III). Without neutralising the gel during bacterial growth, the antimicrobial gel reduced the number of live bacteria in the 3.30, 2.10 and 1.10 μm aerosols by 76%, 69% and 64%, respectively (p = 0.001).

**Figure 3:**
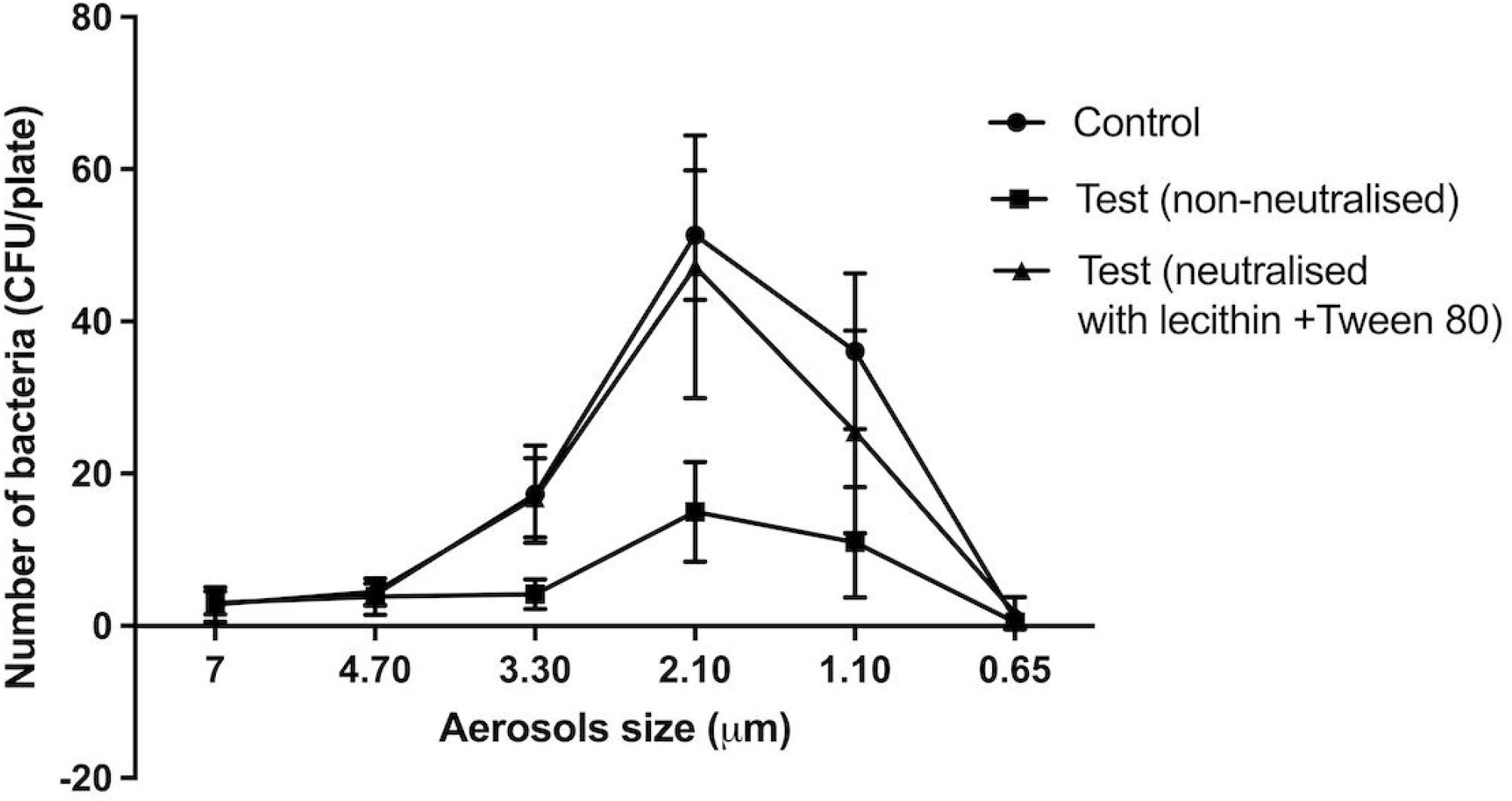
Numbers of *E. coli* K12 recovered from different aerosol sizes with or without neutralisation of the gel vapourised for 10 minutes. The gel significantly reduced the viability of bacteria in aerosols sizes 3.30 and 2.10 μm when neutralised or non-neutralised during bacterial growth compared to untreated control (p < 0.05). Data points represent the mean (± 95% confidence interval) of three independent experiments.

**Table III:** The ability of antimicrobial gel vapourised for 10 minutes to reduce the numbers of aerosolised *E. coli* K12.

| Sample | Amount<br>of gel<br>(g) | Evaporation<br>time<br>(minutes) | Number of<br>bacteria<br>(CFU/mL) | Log<sub>10</sub><br>reduction | %<br>Reduction | p-Value |
| --- | --- | --- | --- | --- | --- | --- |
| Control (no<br>neutraliser) | - | - | 86 ± 14 | - | - | - |
| Antimicrobial gel<br>(No neutraliser) | 10 | 10 | 42 ± 20 | 0.3 | 51.16 | 0.001 |
| Control<br>(With neutraliser) | - | - | 139 ± 46 | - | - | - |
| Antimicrobial gel<br>(With neutraliser) | 10 | 10 | 99 ± 47 | 0.1 | 28.77 | 0.018 |

Similarly, the antimicrobial gel vapours were also active against aerosols *of Aspergillus flavus* spores. The spores mainly travelled in aerosols of between 7.00 μm to 2.10 μm, with significantly (p < 0.001) higher numbers in 2.10 μm than other aerosols sizes (Figure 4). No spores travelled in aerosols of 1.10 μm or 0.65 μm (Figure 4). Overall, the antimicrobial gel reduced the viability of spores of *Aspergillus flavus* by 72% in non-neutralised conditions and 67% when neutralised (p ≤ 0.008; Table IV). Following exposure to the gel, the number of spores in aerosols of 2.10 μm was reduced compared to control by 60% and 73% in neutralised and non-neutralised conditions, respectively (p=0.001). There was no significant difference of the activity if the antimicrobial gel was neutralised or not when growing *Aspergillus flavus* spores (p ≤ 0.08).

**Figure 4.**
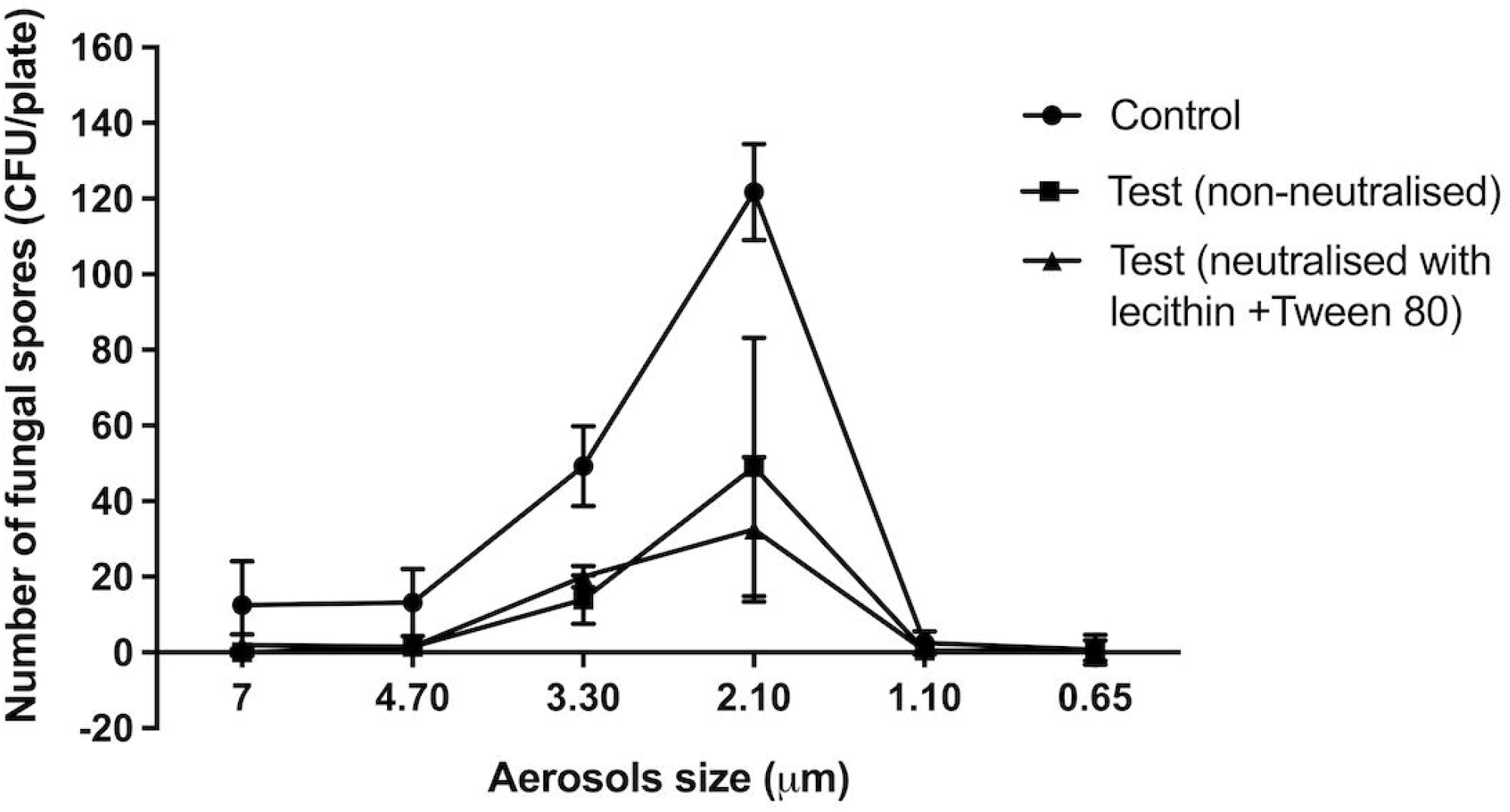
Number of *A. flavus* spores recovered from different aerosol sizes with or without neutralisation of the gel vapourised for 10 minutes. The antimicrobial gel significantly reduced the viability of spores of *A. flavus* in aerosols sizes 3.30 and 2.10 μm in both neutralised and non-neutralised condition compared to untreated control (p < 0.05). Data points represent the mean (± 95% confidence interval) of three independent experiments.

**Table IV:** The ability of the antimicrobial gel vapourised for 10 minutes to reduce the numbers of aerosolised *A. flavus* spores.

| Sample | Amount of<br>gel (g) | Evaporation<br>time<br>(minutes) | Number of<br>fungal<br>spores<br>(CFU/mL) | Log<sub>10</sub><br>Reduction | %<br>Reduction | p-Value |
| --- | --- | --- | --- | --- | --- | --- |
| Control<br>(No neutralizer) | - | - | 231 ± 42 |  |  |  |
| Antimicrobial gel<br>(No neutralizer) | 10 | 10 | 65 ± 7 | 0.6 | 71.86 | 0.008 |
| Control<br>(With neutralizer) | - | - | 170 ± 77 | - | - |  |
| Antimicrobial gel<br>(With neutralizer) | 10 | 10 | 57 ± 23 | 0.5 | 66.47 | 0.001 |

## DISCUSSION

This study has demonstrated that the Mould Gone gel (San-Air) containing compounds found in essential oils of *Melaleuca* genus plants has good antimicrobial activity against bacteria, fungal spores, and coronavirus. Direct contact with the gel resulted in a complete kill of the virus. The gel vapours were able to reduce the numbers of the coronavirus MHV-1 and the bacterium *E. coli* by ≥ 50% and the reduce the number of *A. flavus* spores that can germinate by ≥66%. The ability of vapourised gels to prevent the growth of bacteria, fungal spores and reduce the infectivity of coronavirus in aerosols has not been previously demonstrated, as far as the authors can tell.

This study used a ready-made bacterial filtration efficiency testing rig that is usually used to assess the ability of face masks to filter the bacterium *Staphylococcus aureus* as specified by standard ASTM F2101-1[34]. The Andersen impactor has been widely used to sample environmental bacteria and fungi [35] and has the advantage of being able to directly capture the biological aerosols on agar plates which can then be incubated to directly culture the organisms. Neutralizing chemicals can also be incorporated into the agar to inactivate antimicrobial agents present in the aerosols. While viruses can be captured on the agar plates, they had to be recovered from the agar and cultured on susceptible cells. Other researchers have used this method to culture virus in aerosols [36, 37]. A major advantage of the Andersen impactor is that it allows differentiation of the size of the aerosols in which microbes travel [38].

Human activities such as speaking, coughing and sneezing generate microbial aerosols in sizes ranging from <1 μm to >100 μm [39-42]. Larger aerosols or droplets remain airborne for a short time and settle close to the source [43]. Smaller aerosols under 5 μm in size can remain airborne for longer periods and are able to make their way to the lungs [44]. Aerosols of this size are implicated in the airborne transmission of *Mycobacterium tuberculosis* [45], *Aspergillus fumigatus* spores [46], and viruses including the influenza virus [47] and SARS CoV [48].

The number of viral copies that are needed to cause an infection is expressed as ID_50_ which denotes the mean dose that causes an infection in 50% of susceptible subjects. While the ID_50_ for SARS-CoV-2 is not known, the ID_50_ for SARS CoV ranged from 16 to 160 PFU/person [49]. The vapours produced by the antimicrobial gel were able to significantly reduce the number of viable viral particles in aerosols under 5 μm by 48% within 10 minutes and the reduction increased to 53% when the gel was allowed to vaporise for 20 minutes indicating sustained and perhaps increasing antimicrobial activity. Allowing the gel to vapourize for longer durations and reducing the air flow (i.e., increasing the time for the virus and vapour to interact) may result in further reductions in viral numbers and this should be tested in future experiments.

The gel vapours were bacteriostatic as their activity was diminished in the presence of agents that neutralized their antimicrobial compounds while the viral and fungal activity was unaffected by neutralizers. The active compounds present in the antimicrobial gel used in this study concentrate to approximately 0.0005% (v/v) on evaporation (data supplied by the manufacturer). In-vitro studies performed using essential oils have shown that the active compounds in essential oils are bactericidal for *E. coli* at higher concentrations and bacteriostatic at lower concentrations [50]. Essential oil vapours can affect spore formation in *A. fumigatus* [51] and can be either fungistatic or fungicidal depending on the active compound [52]. Very few studies have examined the antiviral efficacy of essential oil vapours against viruses. One study reported that aerosolised influenza virus or bacteriophage M13 exposed to vapourised essential oils of tea tree or eucalyptus for 24 hours. This resulted in approximately 87% reduction in influenza viral titres, but only 25-42% reduction in M13 titres [27]. Bioactive compounds are present in the essential oils at significantly greater concentration compared to the antimicrobial gel used in this study, and there are concerns about the impact of vapouring these indoor and their impact on human occupants [53]. The low concentration of the active compounds present in the antimicrobial gel are unlikely to impact human health.

This study has demonstrated that vapours produced by a gel containing naturally occurring active compounds can significantly reduce the viable numbers of aerosolized microbes including coronavirus. Using the gel may reduce the transmission of respiratory pathogens, improve indoor air quality and health of human occupants.

